# Ruvbl1 is required for the reproduction of the corn planthopper, *Peregrinus maidis*

**DOI:** 10.1101/2024.04.17.589929

**Authors:** César A. D. Xavier, Clara Tyson, Anna E. Whitfield

## Abstract

*Ruvbl1* (also known as TIP49, Pontin) encodes an ATPase of the AAA+ protein superfamily involved in several cellular functions, including chromatin remodeling, control of transcription, and cellular development (motility, growth, and proliferation). Here, we used an *in-vivo* RNA interference (RNAi) approach to evaluate the effect of *Ruvbl1* silencing on the physiology of the corn planthopper, *Peregrinus maidis*. Silencing of *P. maidis Ruvbl1* (*PmRuvbl1*) was correlated with visible morphology changes in female individuals with significant increases in body mass observed at 8 and 12 days after double strand RNA (dsRNA) injection. Ovary morphology was significantly affected in adult females with *PmRuvbl1* silenced, with no mature oocytes observed at 8 and 12 days after gene silencing. Whereas no significant difference in egg laying was observed 4 days after dsRNA injection, significantly fewer eggs were laid in plants at 8 and 12 days after dsRNA treatment. Furthermore, dramatic reductions in egg hatching were observed at all time points after *PmRuvbl1* silencing, compared to dsGFP-injected controls. These results extend PmRuvbl1 functions as a putative regulator of *P. maidis* reproduction and demonstrate the potential of *Ruvbl1* to be further exploited as a target for RNAi-mediated insect control.

## Introduction

The corn planthopper, *Peregrinus maidis* (Hemiptera: Delphacidae), is a major pest of maize and sorghum in tropical and subtropical areas (Tsai, 2008). In addition to the direct damage due to feeding in the plant vasculature and oviposition in the leaf midribs, this pest indirectly damages plants by transmitting viruses, including maize mosaic virus (MMV) and maize stripe virus (MSpV; Nault and Ammar, 1989, Singh and Seetharama, 2008). Although a MMV resistance gene has been identified in maize germplasm (gene *Mv*; Ming et al., 1997), no resistant genotypes to *P. maidis* have been reported and chemical insecticides are the main strategy used for insect control (Tsai et al., 1990, Higashi et al., 2013). Because of the high reproductive capacity of females, which can lay over 600 eggs during their life span (Tsai, 1996), resistant individuals have a great potential to be selected and replace susceptible insect populations in maize agroecosystems. New technologies to control this pest have been suggested (*e*.*g*. RNA interference, CRISPR-mediated control) and have potential to be exploited as an alternative to chemical control (Yao et al., 2013, Klobasa et al., 2021, Patil et al., 2023, Wang et al., 2024). RNA interference (RNAi) has been extensively used for functional genomics in animals (vertebrates and invertebrates; Bellés, 2010, Podolska and Svoboda, 2011) and plants (McGinnis, 2010), and it has shown great potential to be used for controlling insect pests (Head et al., 2017, Shaffer, 2020, Zhu and Palli, 2020). As a first step, target genes specifically affecting insect physiology (*e*.*g*. survival, development, reproduction) have to be characterized. While RNAi has been developed and demonstrated to be efficient in *P. maidis*, only a few potential target genes have been identified and functionally characterized (Yao et al., 2019, Wang et al., 2023, Wang et al., 2024, Xavier et al., 2024).

The AAA+ protein superfamily is composed of a large number of functionally diverse ATPases with a conserved AAA+ catalytic module that are involved in a tremendous array of cellular processes (Snider and Houry, 2008). AAA+ proteins participate in a variety of functions dependent on energy from ATP hydrolysis for macromolecular remodeling, including DNA replication, ribosome assembly, protein repair, recombination, and beyond (Khan et al., 2022). Ruvbl1 (also known as Tip49, Pontin) is a highly conserved ATPase within the AAA+ protein superfamily with roles in diverse cellular functions, including chromatin remodeling, transcriptional control, cellular development, histone modification, and macromolecular complex assembly (Jha and Dutta, 2009, Dauden et al., 2021). These functions are likely conducted via target gene repression and potentially via DNA helicase activity (Gorynia et al., 2011). It was demonstrated that Ruvbl1 plays essential roles in embryogenesis in mice and *Xenopus laevis* (Bereshchenko et al., 2012, Kim et al., 2023) and its silencing significantly affected cellular development and proliferation impairing larval development in *Drosophila* (Bellosta et al., 2005, Schirling et al., 2010, Prozzillo et al., 2021). However, putative roles of Ruvbl1 in the physiology of non-model insect pests have not been investigated.

Here we evaluated the effect of *Ruvbl1* silencing on the physiology of the corn planthopper, *P. maidis*. Because Ruvbl1 plays multiple regulatory roles in cell function and development, we hypothesized that RNAi-mediated *P. maidis Ruvbl1* (*PmRuvbl1*) silencing would negatively affect insect physiology. The findings presented here provide evidence supporting PmRuvbl1 as a putative regulator of *P. maidis* reproduction.

## Material and methods

### Insect rearing and experimental conditions

An age synchronized *P. maidis* colony has been maintained in the Plant-Virus-Vector Interaction lab at North Carolina State University as previously described (Yao et al., 2019, Xavier et al., 2024). For RNAi experiments, brachypterous females approximately three days after adult emergence from an age-synchronized colony were used. All experiments were conducted under 12:12 light/dark photoperiod at 26°C in a reach-in growth chamber (Conviron), exactly the same conditions used for insect colony rearing.

### Total RNA extraction and reverse transcription quantitative real time PCR (RT-qPCR)

For gene expression analyses, whole-body individuals (first-stage nymph n = 20, second nymph n = 10, third nymph n = 10, fourth nymph n = 5, fifth nymph n = 5, adult males n = 3 and adult females n = 3) and tissues (ovaries n=5 and guts n=6) were pooled prior RNA extraction. Tissue dissection was performed as described previously (Xavier et al., 2024). For RNA extraction, pooled samples were homogenized using tissue Lyser II (Qiagen) in a 1.7 ml microcentrifuge tube containing trizol and five 3 mm glass beads (Pyrex). Total RNA was extracted using Trizol Reagent (ThermoFisher) followed by ethanol precipitation and DNase treatment with TURBO DNA-free Kit (Invitrogen), according to manufacturer’s instructions. The concentration and purity of total RNA was checked with a NanoDrop One spectrophotometer (ThermoFisher) and one microgram of total RNA was used for cDNA synthesis using the Verso cDNA Synthesis Kit (ThermoFisher) in a final volume of 20 μl, following the manufacturer’s instructions.

RT-qPCR reactions consisted of 4 μl of 8-fold diluted cDNA, 1 μl of mixed primers containing 5 μM of each primer and 5 μl of iTaq Universal SYBR Green Supermix (BioRad), as previously described (Xavier et al., 2024). The RT-qPCR was performed in technical duplicates in a CFX Connect Real-Time System (BioRad). Cycles consisted of a denaturation step at 95°C for 1 min, followed by 40 cycles of 95°C for 15 sec and 60°C for 1 min followed by melting curve analysis. *PmRuvbl1* RNA levels were quantified by comparative cycle threshold method (2^-ΔΔCT^) for gene silencing upon dsRNA microinjection and normalized RNA abundance (2^-ΔCT^) was used to evaluate *PmRuvbl1* expression across developmental stages and tissues (Livak and Schmittgen, 2001). *P. maidis ribosomal protein L10* (*PmRPL10*) was used as an internal reference gene (see **Table 1** for primers information; Barandoc-Alviar et al., 2016).

**Table 1.**
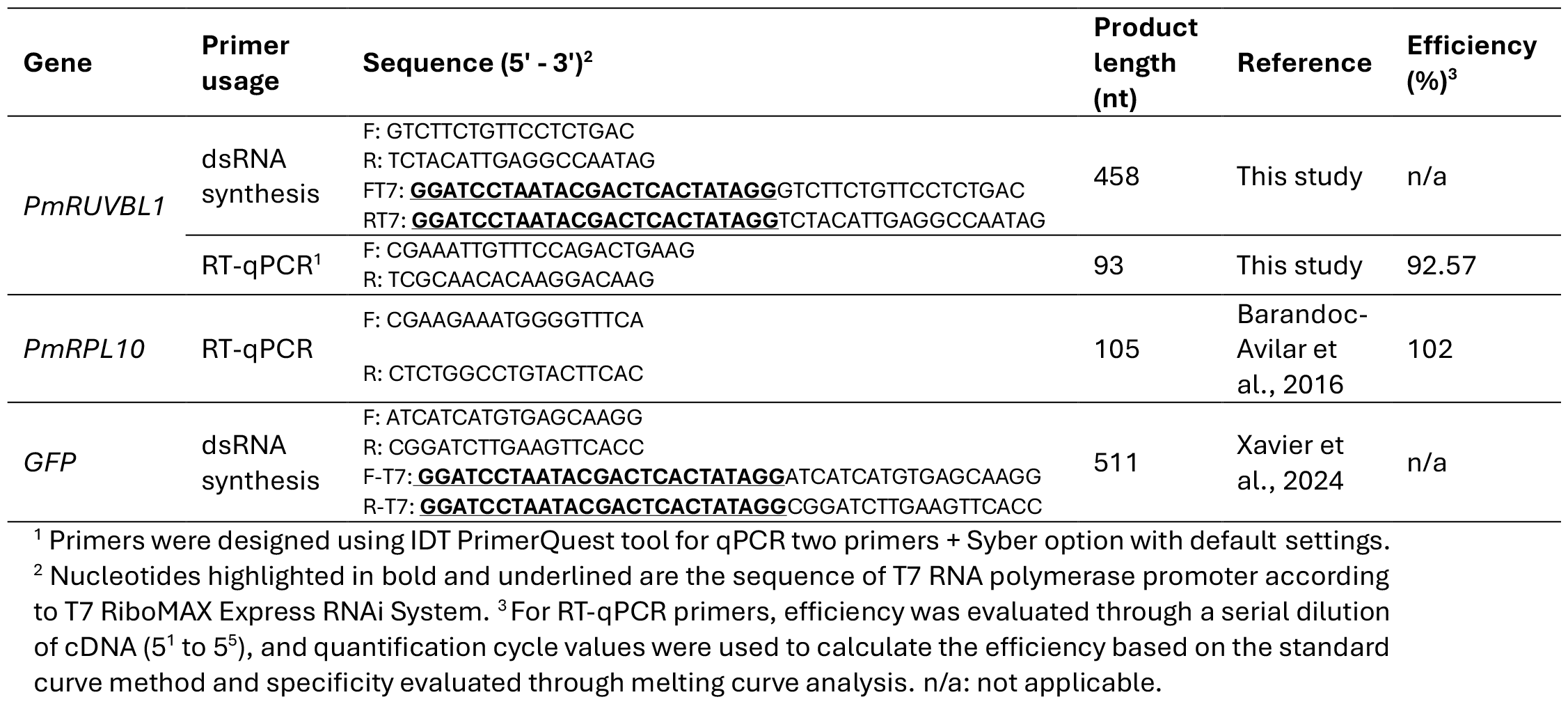
Primers used in this study for RT-qPCR and double strand RNA synthesis are described.

### RNAi assay

RNAi was conducted as previously described by Xavier et al., (2024). Briefly, double-stranded RNA (dsRNA) was synthesized with T7 RiboMAX Express RNAi System (Promega), according to manufacturer’s instructions. Primers were designed following the T7 RiboMAX Express RNAi System guidelines and are described in **Table 1**. DNA template for *in vitro* transcription were obtained by cDNA amplification using Q5 Hot Start High-Fidelity 2X Master Mix (NEB) following manufacturer’s guidelines. PCR products were purified using Monarch PCR & DNA Cleanup Kit (5 μg; NEB) and checked on 1% agarose gel and approximately 600 ng of each PCR product was used as template for *in vitro* dsRNA synthesis according to the manufacturer’s protocol. After synthesis, dsRNA was purified using Monarch RNA Cleanup Kit (50 μg; NEB) and checked on 1% agarose gel and concentration measured with a NanoDrop One spectrophotometer (ThermoFisher). Double-stranded RNA targeting the gene encoding the green fluorescent protein (dsGFP) was used as a negative control.

For dsRNA microinjection, insects were anesthetized on ice and microinjected with 80 nanoliter of 1,000 ng/μl of dsRNA targeting *PmRuvbl1* (dsRUVBL1) or dsGFP at speed of 50 nl/sec using a Nanoinjector III (Drummond Scientific) under a Leica S APO stereo microscope. Insects were injected on the membrane between the meso- and meta-thoracic legs and allowed to recover for one day after microinjection on healthy maize plants before experiments were performed.

### Reproduction and phenotypic analyses

Reproduction assays were performed as previously described (Xavier et al., 2024). Briefly, upon *PmRuvbl1* silencing, the number of eggs laid, and hatched nymphs were counted at time intervals of 4 days during a 12-day period. For oviposition assays, two groups of four females were reared on the 2^nd^ and 3^rd^ oldest leaves of a 2-weeks old maize plant (cv. Early sunglow) using a rectangular clip cage. After each time interval, eggs were visualized and counted under a stereomicroscope following staining with McBride’s solution and destaining with lactic acid:glycerol:water (1:1:1), as previously described (Backus et al., 1988). To quantify the number of eggs hatching, four females were transferred onto a single 7-day old maize plant (cv. Early Sunglow) and given an oviposition period according to each time interval followed by female removal. Plants were allowed to grow, and the number of hatched nymphs was counted. Insect morphology and weight were monitored and evaluated at 4, 8 and 12 days after dsRNA microinjection and imaged with a Leica MC 190 HD camera in a stereomicroscope Leica M205C. Groups of four females were weighed using a precision scale. To verify ovary morphology, females were dissected at 4, 8 and 12 days after dsRNA microinjection under a stereomicroscope in ice-cold 60% ethanol. Photos were taken immediately after dissection with a Leica MC 190 HD camera coupled to the stereomicroscope. These experiments were repeated at least three independent times for each time point tested.

## Results

To investigate putative roles of *PmRuvbl1*-encoded protein in the physiology of *P. maidis*, we first identified its sequence in previously assembled transcriptome data (Martin et al., 2017, Wang et al., 2023). We performed a BLASTn search using *Drosophila melanogaster Ruvbl1* (*DmRuvbl1*, accession number NM144351.4) mRNA sequence as query. Our transcriptome search resulted in two transcripts homologous to *DmRuvbl1*. The sequences shared 100% of nucleotide identity between them and were predicted to encode an open reading frame (ORF) of 1,371 nt, which exactly matches the size of those ORFs encoding Ruvbl1 of *Drosophila* and other planthoppers. Domain analysis using Simple Modular Architecture Research Tool (SMART; Letunic et al., 2020) identified the conserved AAA+ superfamily of ATPases domain (E-value = 2.81^-11^). Further sequence comparison showed that the predicted PmRuvbl1 were highly similar to the putative Ruvbl1 of the planthoppers *Laodelphax striatellus* (97.59% aa identity; accession number RZF35159) and *Nilaparvata lugens* (96.49% aa identity; accession number XP039282926). Furthermore, Ruvbl1 sequences of planthoppers and *Drosophila* shared a high identity at amino acid level (≥80.40% aa identity), suggesting functional conservation.

We next measured RNA levels of *PmRuvbl1* in whole-bodies of nymphs and adults and female tissues (gut and ovary) using RT-qPCR (**Figure 1**). Significant difference in *PmRuvbl1* expression was observed across developmental stages (*F*_6,28_=3.138, *P*=0.0164). Whereas *PmRuvbl1* expression was significantly lower in males compared with females, no significant change in *PmRuvbl1* expression was observed among the five nymphal stages and between nymphs and adults (**Figure 1a**). At tissue level, significantly higher *PmRuvbl1* expression was detected in female guts compared to ovaries (*P*<0.0001; **Figure 1b**). These results demonstrate that *PmRuvbl1* is expressed across all developmental stages and tissues analysed and may indicate variation in biological functions played by PmRuvbl1 across different sex and tissues in *P. maidis*.

**Figure 1.**
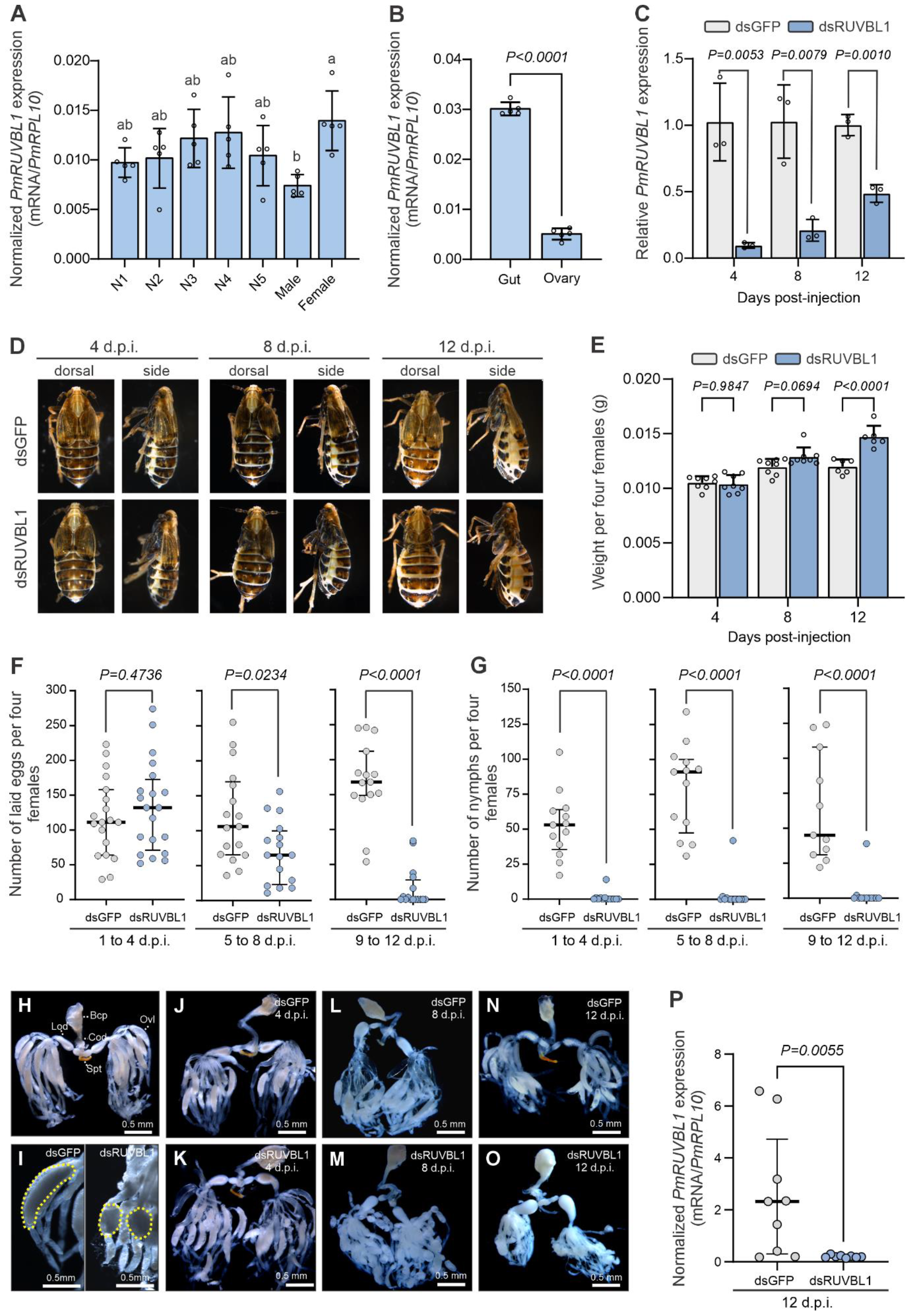
*Peregrinus maidis Ruvbl1* (*PmRuvbl1*) silencing negatively affected ovary morphology and reproduction of females. *PmRuvbl1* expression was analysed through RT-qPCR across (**A**) developmental stages including nymphs (N1 to N5) and adults (male and female), and (**B**) female tissues (gut and ovaries). In **A** and **B** dots represent a biological replication composed of a variable number of pooled individuals according to the stage or tissue (see Method section). In **A**, different letter indicates significant difference among groups as determined by one-way ANOVA followed by Tukey’s multiple comparison test (*P*<0.05). In **B**, significant difference was assessed by unpaired two-tailed t-test (*P*<0.05). Normalized *PmRuvbl1* expression was quantified using 2^−ΔCT^ and *P. maidis ribosomal protein L10* (*PmRPL10*) as an internal reference gene (**A** and **B**). (**C**) *PmRuvbl1* silencing in female whole bodies at 4, 8 and 12 days-post dsRNA injection (d.p.i). Newly emerged females were microinjected with 80 nl of 1,000 ng/μl of dsGFP or dsRUVBL1. Difference across treatments was assessed by unpaired two-tailed t-test (*P*<0.05). Each dot represents a biological replicate composed of three pooled insects prior to RNA extraction. Relative *PmRuvbl1* expression was quantified using 2^−ΔΔCT^ and *PmRPL10* as an internal reference gene. (**D**) Females *Ruvbl1* silenced showed a significant increase in body mass over time. A representative picture is shown. (**E**) Groups of four females were weighted at 4, 8 and 12 d.p.i. Significant differences between groups were assessed by two-way ANOVA followed by Sidak’s test (*P*<0.05). One representative experiment is shown from two independent experiments with similar results. In **A, B, C** and **E**, bars indicate mean and error bars standard deviation. (**F** and **G**) Ruvbl1 silencing led to reduced fecundity of females with reduced oviposition (**F**) and egg hatching (**G**). Error bars represent median and interquartile range of pooled data from three independent experiments. Mann–Whitney test was used to test the statistical significance between groups (*P*<0.05). Each dot represents one biological replicate including the number of eggs laid (**F**) and hatched (**G**) from groups of four females at time interval of 4 days, as indicated in the figure. (**H**-**O**) *PmRuvbl1* silencing led to significant alteration in oocytes morphology. Insects were dissected and pictures of one representative insect from three independent experiments are shown. In **H**, healthy ovary prior dsRNA microinjection is shown. Oocytes became rounded shaped over time in dsRUVBL1-treated females (**I**). (**P**) Efficient *PmRuvl1* was observed in ovaries 12 d.p.i. Each dot represents a single ovary and error bars median and interquartile range. Mann–Whitney test was used to assess the statical significance between groups (*P*<0.05). Ovaries were dissected and RNA extracted using Chelex reagent as previously described (Xavier et al., 2024). Statistical analyses were performed using GraphPad Prism 10.2.1 software. Bcp: bursa copulatrix; Cod: common oviduct; Lod: lateral oviduct, Ovl: ovariole; Spt: spermatheca.

The role of Ruvbl1 in somatic and germ cells has been previously demonstrated and inhibiting its function has been correlated with developmental abnormalities in mammal and *Drosophila* cells (Bellosta et al., 2005, Bereshchenko et al., 2012, Kim et al., 2023). To evaluate the potential of using *Ruvbl1* as a target for RNAi-mediated insect control, we investigated whether *PmRuvbl1* silencing would disrupt female physiology resulting in any significant phenotypic alteration. For that, we used an *in vivo* RNAi approach to silence *PmRuvbl1* through dsRNA microinjection in whole body of newly emerged adult females. Efficient and significant gene silencing was obtained at 4, 8 and 12 days after dsRNA injection with transcript abundance reduced by 91.94%, 81.78% and 51.52%, respectively (*P*<0.05; **Figure 1c**). During the course of silencing experiments, we noticed significant phenotypic alterations in abdomen size and in the number of emerging nymphs from dsRUVBL1-treated individuals compared to control (**Figure 1d-p**). To further investigate whether phenotypic changes were significantly correlated with *PmRuvbl1* silencing, a time course experiment was performed where phenotype and insect reproduction parameters were monitored and recorded. Clear increase in abdomen size was visually observed in dsRUVBL1-treated females which was correlated with significant increase in weight (**Figure 1e, d**). Whereas no significant difference in female weight was observed 4 days after dsRNA treatment (*P*=0.9847), a marginally significant increase was observed at 8 days (*P*=0.0694) with females becoming significantly heavier at 12 days after *PmRuvbl1* silencing (*P*<0.0001), compared to control (**Figure 1e**).

We then investigated the effect of *PmRuvbl1* silencing in female fecundity by evaluating oviposition and egg hatching *in planta*. After dsRNA microinjection, females were let to lay eggs at time intervals of 4 days for a 12-day period. After each time interval, the number of eggs laid, and eggs hatched were evaluated. Whereas no significant difference in egg laying was observed from 1 to 4 days after dsRNA microinjection (*P*=0.4736), significantly fewer eggs were laid in plants from 5 to 8 days (*P*=0.0234) with an even more pronounced reduction observed from 9 to 12 days after dsRNA microinjection (*P*<0.0001; **Figure 1f**). Furthermore, dramatic reduction in egg hatching were observed at all time points after *PmRuvbl1* silencing, compared to dsGFP-microinjected controls (*P*<0.0001; **Figure 1g**). To obtain any clue as to whether reduced fecundity was related to direct effect in ovary functions, we looked at ovary morphology (**Figure 1h-o**). No significant alteration in ovary morphology was observed prior to dsRNA microinjection (**Figure 1h**). Likewise, 4 days after dsRNA microinjection no significant difference in ovaries were observed between dsRUVBL1 silenced and dsGFP-treated control (**Figure 1j, k**). This result is consistent with egg laying assay where no significant difference in the number of eggs was observed at the same time interval (**Figure 1f**). In contrast, ovary morphology was significantly affected in adult females with *PmRuvbl1* silenced at 8 and 12 days after injection. Oocytes were small and round in shape in dsRUVBL1-treated females in contrast with banana-shaped ones observed in control individuals (**Figure 1i; l-o**). Analysis of *PmRuvbl1* expression confirmed that gene silencing was efficiently achieved in ovaries of dsRUVBL1-treated insects (**Figure 1p**). These results extend PmRuvbl1 functions as a possible positive regulator of *P. maidis* reproduction.

## Discussion

Ruvbl1 is an evolutionary conserved protein in eukaryotes from yeast to humans involved in several vital regulatory processes (Jha and Dutta, 2009). Whereas its role in cell physiology and viability has been extensively studied in mammals and *Drosophila* models, little information is available for non-model organisms. Our findings further extend Ruvbl1 functions as a putative regulator of *P. maidis* reproduction. It is well established that Ruvbl1 plays multiple roles involved in cell development and proliferation and modulating its activity by either depletion or overexpression can lead to several cellular developmental abnormalities such as arrested larval development, cellular death, cancer and others (Bellosta et al., 2005, Jha and Dutta, 2009, Wang et al., 2018, Li et al., 2023). Our results suggest that PmRuvbl1 silencing likely affected *P. maidis* fecundity at different stages of the reproductive process. First, at early time points after gene silencing, even though no significant morphological changes were clearly observed in ovaries and oocytes of Ruvbl1-depleted females, eggs laid were not viable. Whereas it is not clear whether pre- or post-embryonic developmental processes were affected, these results indicate that efficient systemic RNAi was achieved resulting in lethality. The Ruvbl1-encoded protein has been shown to be essential for viability in yeast, flies, nematodes and mice (Bellosta et al., 2005, Bereshchenko et al., 2012). Strong silencing of Tip60 complex, which includes Ruvbl1, caused pupal lethality in *Drosophila* (Schirling et al., 2010). Furthermore, previous studies demonstrated that loss of function of Ruvbl1 significantly affected early embryogenesis in mice and *X. laevis* (Bereshchenko et al., 2012, Kim et al., 2023). While the mechanism(s) by which Ruvbl1 regulates embryogenesis is still not well understood, the results obtained here and elsewhere reinforce the conserved vital role of Ruvbl1 in early developmental processes involved in reproduction.

At later time points, the silencing of *PmRuvbl1* led to significant morphological alterations of ovarioles and oocytes, with no mature eggs being observed. Basic cellular processes regulated by Ruvbl1 involved in early organism development and growth (*e*.*g*. cell-cycle progression, transcription, cell proliferation, DNA repair) have been well characterized. Therefore, it’s not unexpected that its silencing may affect pre-zygotic events such as oogenesis. Over time, PmRuvbl1 silencing led to smaller and rounded shaped oocytes which is consistent with regulatory roles of Ruvbl1 in cellular growth and proliferation (Bellosta et al., 2005). In *Drosophila*, regulation of cell growth during normal development is mediated by Ruvbl1 interaction with dMyc, the *Drosophila* ortholog of the vertebrate c-Myc, a central regulator of growth and cell-cycle progression (Bellosta et al., 2005). Because of Ruvbl1 multifunctionality, acting as a master regulator and partnering with several other factors related to basic cellular functions (e.g. c-Myc, c-Jun N-terminal kinase), *Ruvbl1* silencing could lead to multiple peripheral effect in the organism physiology. For example, Ruvbl1 is a crucial modulator of c-Jun N-terminal kinase (JNK) pathway during *Drosophila* development and its silencing has been associated with cell-death via JNK pathway (Wang et al., 2018). Modulation of JNK pathway significantly impaired ovary development in swimming crab (Wei et al., 2020). Therefore, silencing hub genes such Ruvbl1 can result in substantial addictive effects negatively affecting cell physiology. Unveiling these interactions and mechanisms may offer additional possibilities for finding targets to be manipulated for insect control.

## Conclusion

Ruvbl1, a highly conserved ATPase, engages in divergent cellular processes; here we demonstrate the effects of RNAi-mediated knockdown of Ruvbl1 on insect morphology and reproduction. Since its discovery in 1998, RNAi technology has transformed entomological research and additionally serves as a promising, highly species-specific alternative to the use of chemical insecticides. Despite widespread use of traditional pesticides, insect pests cause 18 to 20% of annual global crop loss (Sharma et al., 2017), evidencing the need for novel methods of insect pest management. As a reproductive regulator in *P. maidis*, Ruvbl1 represents a putative target gene for population suppression, although additional studies should be performed to address RNAi delivery and evaluate potential effects in non-target organisms.

## Acknowledgement

We thank the members of the Plant-Virus-Vector Interactions Lab team for the valuable discussion throughout the course of the project. This project was supported by the USDA Biotechnology Risk Assessment Grants (BRAG) program Grant # 2022-33522-37745 and The North Carolina State University Department of Entomology and Plant Pathology was part of a team supporting DARPA’s Insect Allies Program.

## Notes

### Competing Interest Statement

The authors have declared no competing interest.

